# Liver-specific Deletion Of Small Heterodimer Partner Alters Enterohepatic Bile Acid Levels And Promotes Bile Acid-Mediated Proliferation In Male Mice

**DOI:** 10.1101/2021.09.07.459308

**Authors:** Ryan Philip Henry Shaw, Peter Kolyvas, Nathanlown Dang, Angela Hyon, Sayeepriyadarshini Anakk

## Abstract

Small heterodimer partner (*Shp*) regulates several metabolic processes, including bile acid levels, but lacks the conserved DNA binding domain. Phylogenetic analysis revealed conserved genetic evolution of *Shp, Fxr, Cyp7a1* and *Cyp27a1,* underscoring the importance of these molecules in maintaining bile acid homeostasis. *Shp*, although primarily studied as a downstream target of Farnesoid X Receptor (*Fxr*), has a distinct hepatic role that is poorly understood. Here we report that liver-specific *Shp* knockout (*LShpKO)* mice have impaired negative feedback of *Cyp7a1* and *Cyp8b1* upon bile acid challenge and demonstrate that a single copy of the *Shp* gene is sufficient to maintain this response. *LShpKO* mice also exhibit elevated total bile acid pool with higher bile acid fraction in the intestine mimicking the 1% cholic acid (CA) fed control mice. Agonistic activation of *Fxr* (GW4064) in the *LShpKO* did not alter the elevated basal expression of *Cyp8b1* but lowered *Cyp7a1* expression. We found that deletion of *Shp* led to an enrichment of distinct motifs and pathways associated with circadian rhythm, amino and carboxylic acid metabolism, copper ion transport, and DNA synthesis. *LShpKO* livers displayed a higher basal proliferation that was exacerbated specifically with bile acid challenge but not with another liver mitogen, TCPOBOP (TC). Overall, our data indicate that hepatic SHP uniquely regulates certain proliferative and metabolic cues.

## INTRODUCTION

Cholestasis, a condition wherein bile acids (BA) homeostasis is impaired, leading to excess hepatic accumulation, is commonly observed in several liver diseases (1–4). Sustained exposure to BAs induces injury, dysregulation of the cell cycle, apoptosis, and consequently malignant transformation. Thus, it is essential to understand how bile acid levels are controlled.

Nuclear receptors (NR) are ligand-activated transcription factors that coordinate major cellular pathways, including metabolism (5–8). The atypical NR, small heterodimer partner (*Shp*), modulates pathways involved in bile acid homeostasis, fibrolamellar carcinomas, and non-alcoholic fatty liver disease (NAFLD) (9–12). Most studies have been performed using global *Shp*KO mice and because BAs are endogenous ligands that bind Farnesoid X Receptor (*Fxr*) (which in turn can induce *Shp),* the focus has been laid on elucidating FXR function. SHP is recruited to suppress *Cyp7a1* transcription (13–15) to orchestrate a negative feedback loop to prevent BA excess. The severity of the cholestasis in *Fxr/Shp* double knockout mice compared to the individual knockouts revealed cooperative and distinct roles for FXR and SHP (16).

Intriguingly, conditional knockout of *Fxr* in the liver did not increase serum BAs, and *Cyp7a1* and *8b1* suppression in response to *Fxr* agonist GW4064 was maintained. Further, tissue-specific roles for *Fxr*-mediated BA homeostasis have been uncovered with conditional knockout of *Fxr* in the intestine, losing GW4064-mediated suppression of *Cyp7a1* but not *Cyp8b1* gene expression (17–18). Therefore, we investigated the liver-specific *Shp* knockout mice (*LShpKO*) and examined their response to an acute bile acid challenge. We examined the expression of genes responsible for hepatic synthesis, transportation, and enterohepatic BA axis in the absence of hepatic *Shp*.

Mature hepatocytes are quiescent but can be coaxed to proliferate in response to injury. BAs can also act as mitogens, which goes hand in hand with *Shp’s* postulated role in inhibiting liver proliferation (10) (19–31). We examined if deletion of *Shp* altered proliferation after treatment with either CA or ligand-based *Car* activation. Parallelly, we mined the publically available data for *Shp*KO and *Fxr*KO mice and examined unique and overlapping gene sets (Supplemental Tables 1 & 2) (32) to identify distinct pathways that are regulated by these two receptors. Then using computational tools, we examined if there was a common motif of regulation of genes that were uniquely altered when *Shp* was deleted. Finally, to test the relevance of *Shp* upon *Fxr* activation, we treated *LShpKO* mice with GW4064 and examined its response in the liver.

## MATERIALS AND METHODS

### Generation of floxed *Shp* and Liver Specific *Shp* knockout mice

*Ff Shp* mice were obtained from Dr. Kristina Schoonjans’ laboratory. By breeding *Albumin-Cre* mice with *ff Shp* mice we generated heterozygous *Shp* +/-, and homozygous *Shp* -/- liver-specific *Shp* Knock-out (*LShpKO*) mice. *ff Shp* mice were used as controls.

### Animal Experiments

Adult male mice aged 9-21 weeks old were used in this study. Mice were maintained in flow cages at 24°C on the standard 12hr day/night schedule. Food (including experimental diets) and water were available *ad libitum*. Control mice were fed standard chow diet, while cholic acid (CA) experimental group were fed 1% cholic acid for 3 days. Mice were sacrificed via cervical dislocation in the fed state at 9-10 am. Mice were bled Retro-orbitally prior to sacrifice, and serum was extracted after centrifugation at 7,000 rpm for 10 minutes. Serum was transferred to black tubes and stored in -80°C. The liver, gallbladder, and small intestine (ileum) were collected, flash frozen in liquid nitrogen, and stored at -80°C. Mice were cared for according to the National institute of Health (NIH) guidelines and the Institutional Animal Care and Use Committee at the University of Illinois at Urbana-Champaign approved all experimentation. Mice were gavaged with either vehicle (1% Methylcellulose 1% TritonX-100 in PBS) or 50mg/kg *Fxr* agonist GW4064. Mice were gavaged twice, first in the evening, second in the morning and sacrificed 3 hours post second gavage.

### Genotyping Assay

Tail samples were digested in a 200:2 ratio of Tail lysate (VIAGEN 102-T) to proteinase K overnight in 55°C water bath overnight and then inactivated at 85°C for 45 min. 10µl of GoTaq buffer, 0.1µl each of 100µM Forward and Reverse Primers, 7.3µl of water, and 2.5µl of digested tail were used in the *ff Shp* PCR reaction. *Alb-Cre* reaction used 10µl of GoTaq buffer, 1µl of a 10µM dilution mix of Forward and Reverse Primers, 6.5µl of water, and 2.5µl of digested tail. The PCR products were then run on a 1% Agarose gel. WT *Shp* band is seen at 365 bp, whereas floxed *Shp* band is seen at 415 bp. Positive Alb-Cre was observed at 390 bp. Each PCR product was run alongside a 100bp DNA ladder (BioLabs) compared with positive and negative controls.

### Histological Preparation

Liver and Ileum tissue were preserved in 10% neutral buffered formalin for 24 hrs at 4°C, transferred to 75% ethanol and stored until processing (up to 4 weeks). Tissue processing occurred in the following steps with each step occurring for 45 minutes. 95% ethanol 2x, 100% ethanol 3x, and xylene 2x. Samples were then embedded in paraffin blocks. Blocks were sectioned at 5µm and placed on glass slides.

### Hematoxylin and Eosin Staining

Tissue sections were deparaffinized and rehydrated through 3 changes of xylene at 5 minutes, 100% ethanol 3x for 3 minutes, 95% ethanol for 3 minutes, 80% ethanol for 3 minutes, 50% ethanol for 3 minutes, and rinsed in tap water. The slides were then placed in Hematoxylin 7211 (Thermo Fisher) for 1 minute and 50 seconds and quickly rinsed in DI water until clear. Sections were placed in bluing reagent NaHCO_3_ for 1 min, rinsed in DI water, and incubated in Eosin Y (Thermo Scientific) for 25 seconds. Sections were then placed in 100% ethanol 3x for 3 minutes, and xylene 3x for 5 minutes. Sections were left in xylene overnight, and subsequently mounted and coverslipped using Permount (Fisher).

### ALT/AST Serum Analysis

Previously collected serum was thawed on ice and ALT/AST Assay was run according to the Thermo Scientific kit protocol. Absorbances were obtained using a Biotek Synergy 2 reader.

### Immunohistochemistry

Liver samples were preserved, sectioned, deparaffinized, and rehydrated as described above. Tris-EDTA (10mM Tris, 1mM EDTA, .05% Tween 20, pH 9) buffer was used for antigen retrieval, with samples being microwaved for 35 minutes. Slides were cooled and endogenous peroxidases were quenched by incubating slides in 3% H_2_O_2_ in methanol for 15 minutes. Samples were blocked with a buffer of 2% NGS, 1%BSA, .25% Tween 20, .05% Triton X100 in 1XTBS for 1 hr at RT. Samples were then incubated in Primary antibody (Purified mouse Anti Ki-67, BD Biosciences, Cat no: 550609) diluted to 1:100 in dilution buffer (1%BSA, .05% Triton X100, in 1XTBS) overnight at 4°C. Samples were washed in 1X TBST and then incubated in secondary antibody (Goat Anti-Mouse IgG (H+L) HRP Conjugated, BioRad Cat No: 170-6516) diluted 1:250 in dilution buffer at RT for 1 hour. Samples were washed and DAB reagent (Vector Cat no: SK-4100) reagent was added according to manufacturer protocol. Samples were counterstained in Hematoxylin 7211 (Thermo Fisher) for 15 seconds, rinsed in water until it runs clear, placed in bluing reagent NaHCO_3_ for 1 min, and finally rinsed in DI water again. Samples were dehydrated and coverslipped as described above. Ki-67 cell count analyses were performed by taking 5 random images per sample.

### RNA Isolation, cDNA synthesis, and Quantitative Real Time-PCR

The total RNA was extracted from flash-frozen liver and intestinal tissue according to TRIzol (Ambion) manufacturer protocol. RNA quality was checked by measuring the A260/A280 using the Biotek Synergy 2 as well as by 1% Bleach agarose TAE gel analysis. A 5µg volume of RNA was used and treated with DNase I (BioLabs New England) and subsequently deactivated with EDTA and heat. cDNA was made using Maxima Reverse Transcriptase (Thermo Scientific), random primers (BioLabs New England), dNTP mix 10mM (Invitrogen), .1M DTT(BioLabs New England), and 5x RT buffer(Thermo Scientific). cDNA was then diluted to 12.5ng/µl using molecular grade water. qRT-PCR was performed using SYBR green-based reaction at 50ng of cDNA per reaction. Relative expression was determined by using 2^ΔCT, with the control gene being *36B4*. All primer sequences can be found in supplemental table 5.

### Bile Acid Analysis

Total bile acid levels were determined from liver, ileum, gallbladder, and serum using Diazyme’s Total Bile Acid Assay kit. Liver and intestinal tissue was homogenized in 75% ethanol, incubated at 50°C for 2 hrs, and centrifuged at 12,000 rpm for 10 minutes. The supernatant was then diluted 4-5x or 35-40x respectively prior to using the assay. Gallbladders were thawed on ice and then pierced to expel its contents. 1µl of the extract was diluted 20-30x before use in the assay. Serum was thawed on ice and either directly used in assay or diluted 2x. All samples were diluted in 0.9% saline. Samples were aliquoted in triplicates, and obtained absorbance values were compared to a standard curve comprised from 50µmole/L calibrator dilutions. Total bile acid levels were obtained by calculating bile acid concentration in µM/g of tissue used with the total liver weight used as the tissue weight for serum and gallbladder. All concentrations for each tissue were averaged and then summed together to get an average total bile acid level for each group. The colorimetric analysis was obtained via Biotek Synergy 2 instrument.

### Transcription Factor Analysis

Using publically available transcriptome data of *Fxr*KO and *Shp*KO mice, genes that exhibited more than 2 fold change in the knockout compared to control were examined. Genes unique to each of the two genotypes were considered for analysis. We wanted to explore the regulatory sequence of these unique genes that altered only when *Shp* or only when *Fxr* is deleted. To do this, we scanned 600 nucleotides upstream of each of these genes from the transcriptome data of *Fxr*KO and *Shp*KO mice using the Ensembl-BioMart tool and inserted them into MEME-Suite’s Analysis of Motif Enrichment (AME) tool, an in-silico identifier of motifs enriched in the gene promoters. The database of motifs used by AME for this analysis was the HOmo sapiens COmprehensive Model COllection (HOCOMOCO) v11 which derives its mouse motif models from the ChIPMunk motif discovery tool. A motif list for *Fxr*KO and *Shp*KO was made and compared for transcription factor motifs unique to the *Shp*KO genotype. This led to a final list of transcription factors that are possibly uniquely regulated by *Shp* as shown in Supplemental Table 3 and a list of uniquely regulated transcription factors by *Fxr* as shown in Supplemental Table 4 (32). Literature indicates these transcription factors are involved in proliferation and cell cycle progression. As such, genes involved in the E2F cell-cycle progression pathway and a few genes from the transcriptome data of *Fxr*KO and *Shp*KO mice that are cell-cycle associated were analyzed using qPCR. The primer sequences used for their analysis are shown in supplemental table 5.

### TCPBOP Preparation and Injection

1,4- bis[2-(3,5-dichloropyridyloxy)]benzene (TC) was purchased from Sigma-Aldrich in 5mg quantities. 1-1.5 ml of 100% ethanol was added to resuspend the compound, and then 2ml of corn oil was added to the mixture to achieve a 2.5mg/ml concentration. This mixture was left to stir overnight to evaporate any residual ethanol in the solution. The ethanol-free TC and corn oil mixture was then injected intraperitonally into *ff Shp* and *LShpKO* mice. Mice were injected with 5ul of solution per gram weight and then sacrificed 3 days post injection.

### Phylogenetic analysis

The evolutionary history was inferred using the Neighbor-Joining method. The bootstrap consensus tree inferred from 500 replicates is taken to represent the evolutionary history of the taxa analyzed. Branches corresponding to partitions reproduced in less than 50% bootstrap replicates are collapsed. The evolutionary distances were computed using the Poisson correction method and are in the units of the number of amino acid substitutions per site. This analysis involved 77 amino acid sequences. All ambiguous positions were removed for each sequence pair (pairwise deletion option). There were a total of 898 positions in the final dataset. Evolutionary analyses were conducted in MEGA X (33–36).

### Statistical Analysis

All data is represented as mean ± SD. A two-tailed unpaired t-test was used to determine significance between 2 groups. Two-way ANOVA analysis of variance Bonferroni test was used to determine significance between 2 groups under 2 conditions. Statistics were performed using the GraphPad Prism 6 software. Outliers were determined using Graphpad and removed from the analysis. Statistical significance was considered as *P≤.05, **P≤.01, ***P≤.001, ****P≤.0001.

## Supplemental Data

Supplemental figures and tables can be found in an online repository. (32)

## RESULTS

### A single copy of the hepatic *Shp* allele is sufficient to maintain BA homeostasis

We generated liver-specific *Shp* heteroz*ygous (Shp +/- )* and homozygous knockout *(LShpKO) mice* using albumin-specific Cre-recombinase. Both of these genotypes maintained intact expression of the *Fxr* transcript (Supplemental Figure 1 C) (32). Intriguingly, *Shp* expression was comparable between the control mice expressing both functional alleles to that of the heterozygous *Shp* knockout possessing only one functional allele. Next, we challenged *ff Shp, Shp +/-* and *LShpKO* mice with either chow or 1% CA diet and analyzed the expression of genes involved in the BA synthesis and transportation pathways. We found that basal *Cyp7a1* and *Cyp8b1* mRNA levels were significantly higher when *Shp* was deleted in the liver, even under chow conditions (Figure 1 A). The expression of alternative BA synthesis genes, *Cyp27a1* and *Cyp7b1,* remained unchanged irrespective of *Shp* expression. We examined the canalicular and peripheral hepatic export pumps and found that transcript levels of canalicular exporters *Mdr2* and *Mrp4* were increased in *LShpKO l*ivers (Figure 1 B). Upon challenging with a 1% CA diet, *ff Shp* mice showed the expected suppression of BA synthesis pathways *Cyp7a1* and *Cyp8b1*, which was blunted in the absence of hepatic *Shp* (Figure 1 A). However, *Shp +/-* mice maintained this repression of BA synthesis and had comparable expression patterns of the BA transporters as that of *ff Shp* mice, indicating a single copy of the allele is enough to maintain the repressive function (Supplemental Figures 2 & 3) (32). We also observed induction of *Mrp3* mRNA levels but not the other transporters in CA-fed *LShpKO* mice (Figure 1 B).

**Figure 1:**
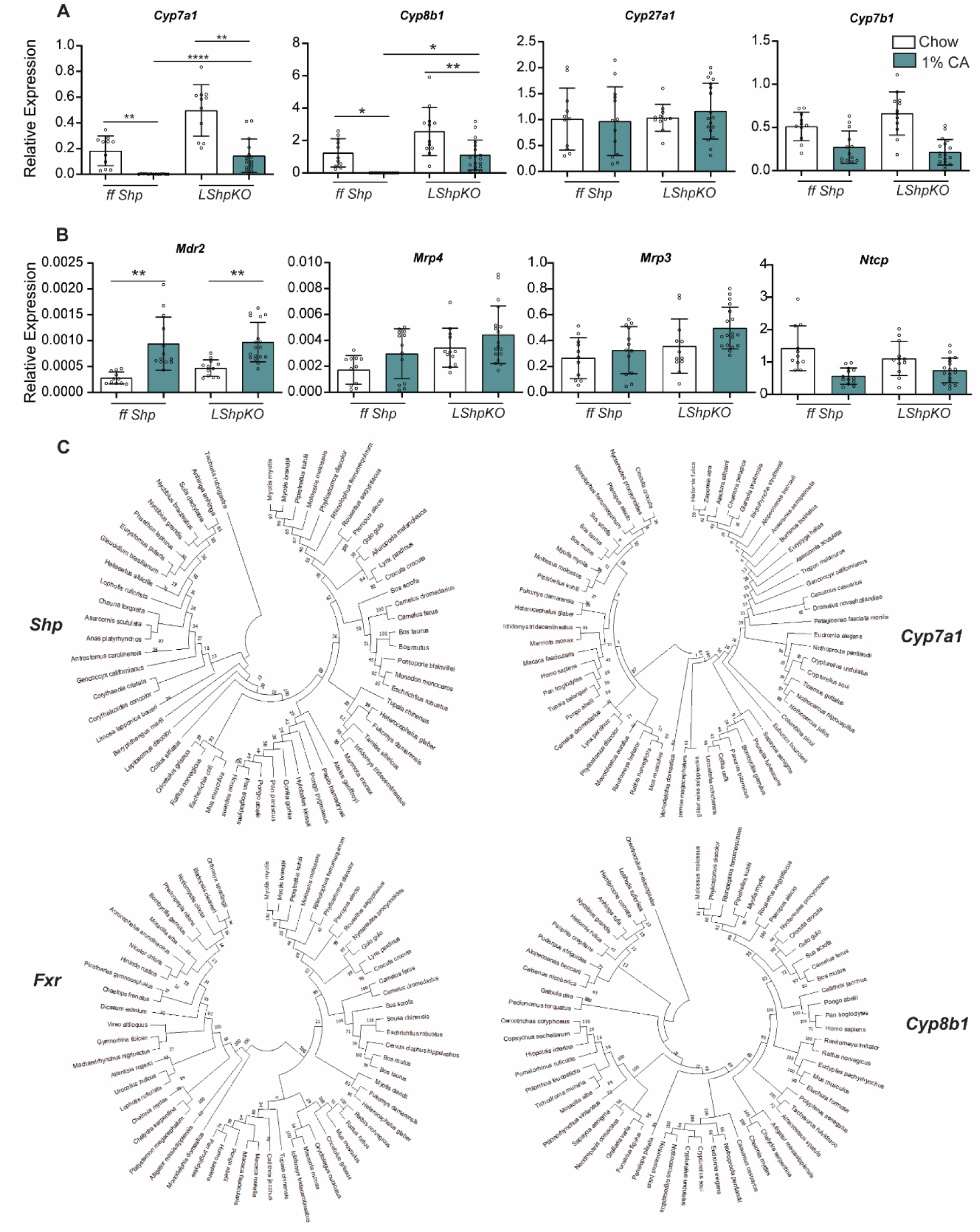
Relative Expression of bile acid synthesis and transport genes in ff *Shp* and *LShpKO* mice on chow and 1% CA diets. (A) Expression of key bile acid synthesis genes in ff *Shp* and *LShpKO* animals under chow and 1%CA conditions. (B) Expression of hepatic bile acid uptake and canalicular bile acid export pumps in ff *Shp* and *LShpKO* animals under chow and 1%CA conditions. (C) Phylogenetic analysis of the evolution of *Shp* from *Fxr, Cyp7a1 and Cyp8b1*. The evolutionary history was inferred using the Neighbor-Joining method. The bootstrap consensus tree inferred from 500 replicates is taken to represent the evolutionary history of the taxa analyzed. Branches corresponding to partitions reproduced in less than 50% bootstrap replicates are collapsed. The evolutionary distances were computed using the Poisson correction method and are in the units of the number of amino acid substitutions per site. This analysis involved 60 amino acid sequences per gene examined. All ambiguous positions were removed for each sequence pair (pairwise deletion option). There were a total of 898 positions in the final dataset. Evolutionary analyses were conducted in MEGA X (43–46)One-way ANOVA analysis of variance Bonferroni test was used to determine significance between groups. *P≤.05, **P≤.01, ***P≤.001, ****P≤.0001.

We also investigated the evolution of SHP-mediated BA regulation by performing phylogenetic analysis using the protein sequences of FXR, SHP, CYP7A1 *and* CYP8B1. We found similarities in the branching structures of FXR and CYP7A1 while the sequences of SHP and CYP8B1 more so resembled each other (Figure 1 C). Since *Shp* is a gene that contains a single intron with two exons, we suspected that was an ancient gene. The phylogenetic analysis suggests that *Shp* is not an ancestral gene, in fact, the two major branches present in the FXR phylogenetic tree are drastically reduced in size in the tree constructed for SHP.

### Loss of hepatic *Shp* leads to elevated BA pool and alters its tissue distribution

To characterize BA homeostasis in *LShpKO* animals, we measured bile acid levels from chow and 1% CA fed control and knockout mice. We found that hepatic *Shp* deficiency shifted the distribution of BAs in enterohepatic tissues (liver, intestine, and gall bladder). Under chow conditions, *LShpKO* animals exhibited higher ileal but lower hepatic BA content than *ff Shp* mice on a chow diet (Figure 2 A). When challenged with CA diet, *LShpKO* animals exhibit a significant decrease in the gallbladder (biliary) content with a stark increase in ileal, liver, and serum BA content compared to *ff Shp* animals (Figure 2 B). In addition to the changes in the distribution of the BA pool, *LShpKO* mice exhibit an overall increase in BA levels than *ff Shp* animals (Figure 2 A & B). In control animals, the biliary bile acid levels are the highest, followed by the ileal content. Upon 1% CA feeding, the ileal BA concentrations increase significantly. *LShpKO* animals under chow diet closely resemble control animals on a 1% CA diet and their ileal bile acid levels are consistently higher than in *ff Shp* animals under both chow and 1%CA diet (Figure 2 C). We investigated several intestinal transport genes to provide insight into why ileal bile acid contributions were elevated in the absence of hepatic *Shp*. Transcript expression of Ileal import and efflux transporters remained similar between *LShpKO* and control mice, and we do not know why this increase in BAs is observed in the intestine. But we observed increased expression of *Fgf15* in *LShpKO* animals under chow conditions. (Supplemental Figure 4) (32).

**Figure 2:**
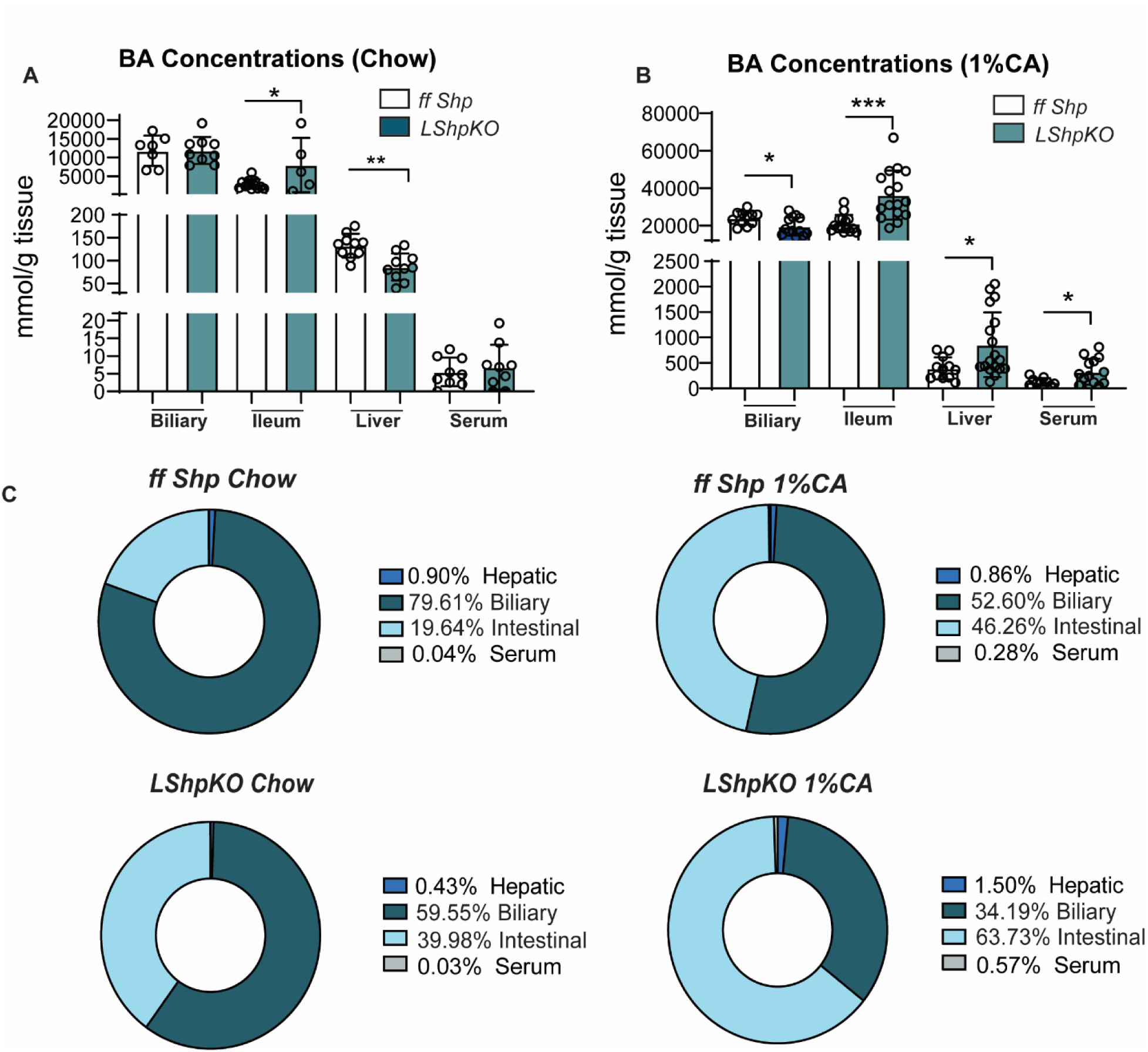
(A) Tissue-specific bile acid levels under chow diet. B) Tissue-specific bile acid levels under 1%CA diet. (C) Relative percent composition of total bile acid pool accounted for by each tissue. N=5-18 mice per group for bile acid assay. One-way ANOVA analysis of variance Bonferroni test was used to determine significance between groups. *P≤.05, **P≤.01, ***P≤.001, ****P≤.0001.

### Loss of hepatic *Shp* induces a mild proliferative response in the liver

We examined liver sections stained with hematoxylin and eosin for inflammatory and morphological changes. *LShpKO* displayed no overt morphological difference; however, under 1% CA conditions, these mice showed increased infiltration of inflammatory cells and occasional necrosis in approximately 30% of livers sampled compared to control animals (Figure 3 A). Hepatic serum injury markers ALT and AST were comparable between *ff Shp* and *LShpKO* animals (Figure 3 B & C).

**Figure 3:**
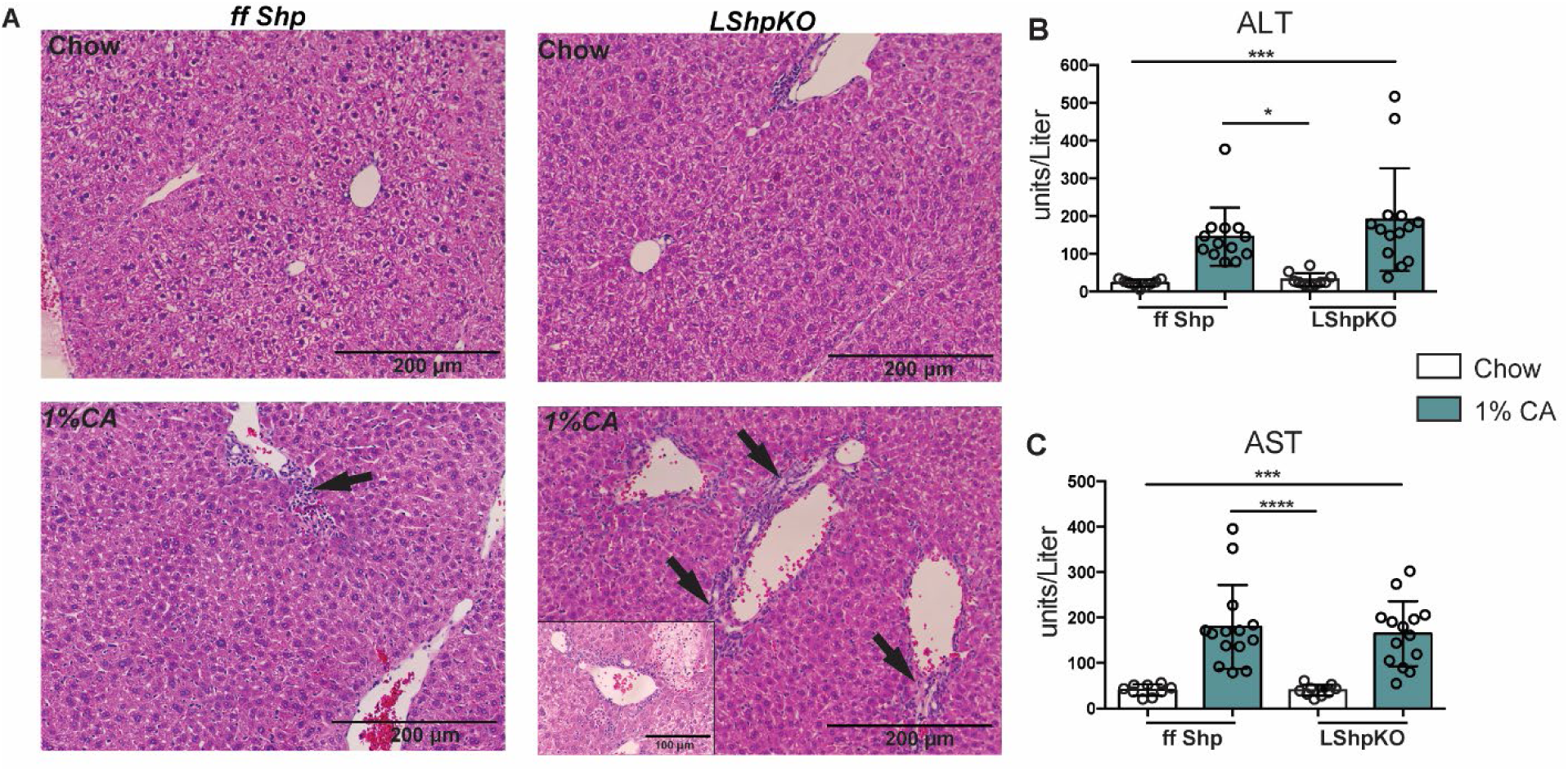
Hematoxylin & Eosin staining of liver tissue and Injury markers. A.) H&E stain with inflammation and damage noted with arrows. B-C.) Liver injury markers ALT/AST levels. N=5-15 mice per group for H&E stainin. N=9-14 mice per group for ALT AST Assay. Values shown as mean ± SD. One-way ANOVA analysis of variance Bonferroni test was used to determine significance between groups. *P≤.05, **P≤.01, ***P≤.001, ****P≤.0001.

To test if the injury secondary to CA diet induced proliferation in mice that lack hepatic *Shp*, we performed Ki-67 immunostaining and quantified the number of hepatocytes and non-parenchymal cells (NPCs) that were undergoing cell division and found more Ki-67 positive nuclei in *LShpKO* animals compared to control animals on a chow diet (Figure 4 A-C). This increase is further exacerbated by 1% CA feeding in *LShpKO* animals.

**Figure 4:**
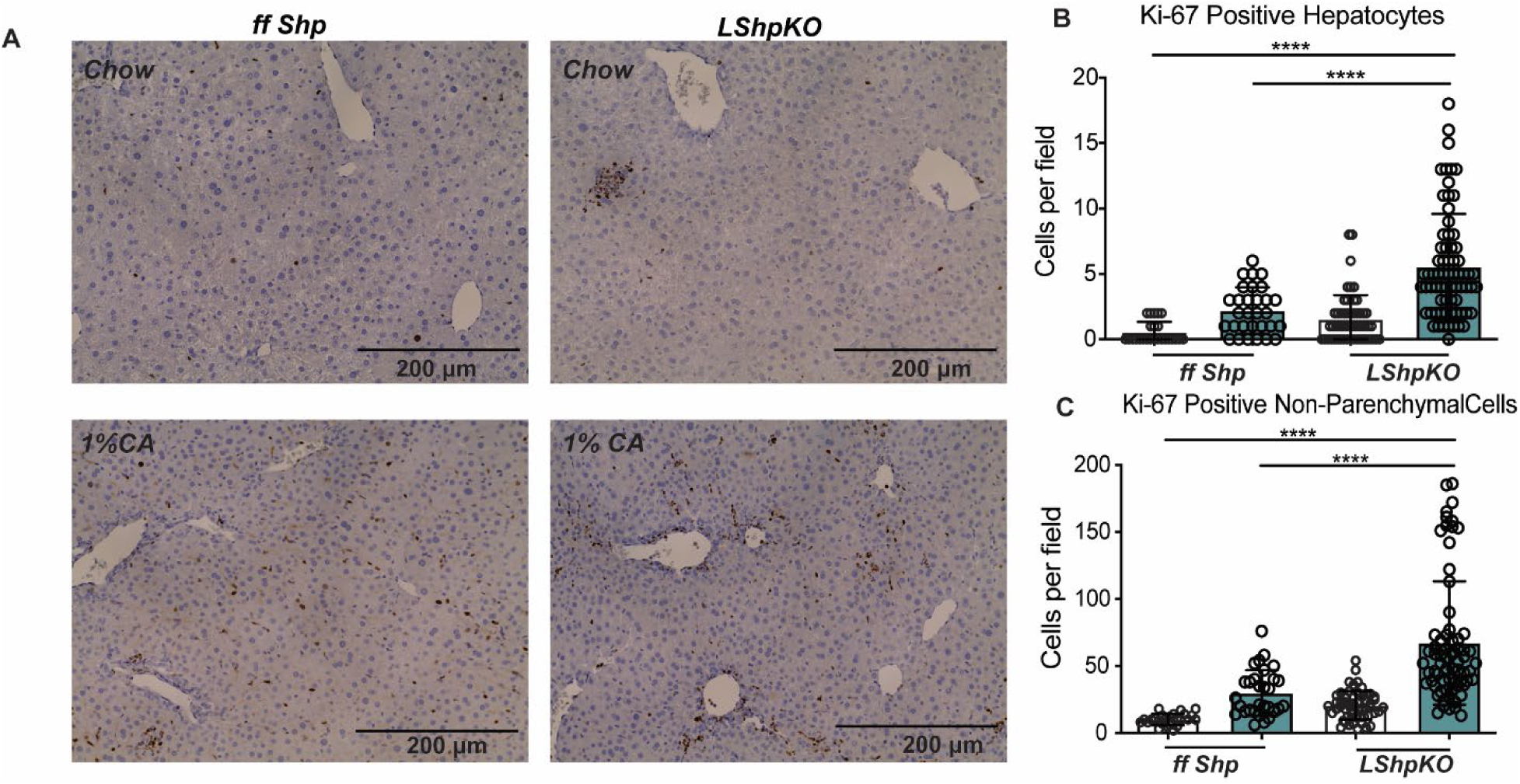
Ki-67 immunostaining and proliferation quantification A.) Ki-67 immunohistochemistry of liver tissue samples showing proliferation of hepatocytes and Non-Parenchymal Cells (NPCs). B.) quantification of proliferating hepatocytes C.) quantification of proliferating NPCs. N=5-14 mice per group with 5 random images taken per mouse. Values shown as mean ± SD. One-way ANOVA analysis of variance Bonferroni test was used to determine significance between groups. *P≤.05, **P≤.01, ***P≤.001, ****P≤.0001.

To test if the excessive proliferative response was unique to BA overload, we challenged *LShpKO* mice with a potent mitogen, TCPBOP (TC) an agonist of another nuclear receptor, constitutive androstane receptor (CAR) (37–40). Upon activation, CAR translocates to the nucleus to induce proliferation and detoxification networks (41). *LShpKO* and *ff Shp* mice were injected with TC or corn oil as a vehicle and we examined the gene expression of several cyclins. TC-mediated *Ccnd1* increase was lost, while *Ccnb2* induction was blunted in *LShpKO* mice (Figure 5 C), suggesting that *Shp* may play a role in the S to the G2 transition in proliferating hepatocytes. On the other hand, increases in *Ccnb1* and *Ccne* transcripts were maintained in the absence of *Shp* (Supplemental Figure 5 C) (32). We then investigated the proliferative changes with immunohistochemistry for Ki-67. Surprisingly, despite a loss in *Ccnd1* and a significant reduction in *Ccnb2* expression, Ki-67 positive nuclei remained comparable with and without the expression of hepatic *Shp,* indicating the presence of compensatory signals in *LShpKO* mice (Supplemental Figure 5 A & B) (32).

**Figure 5:**
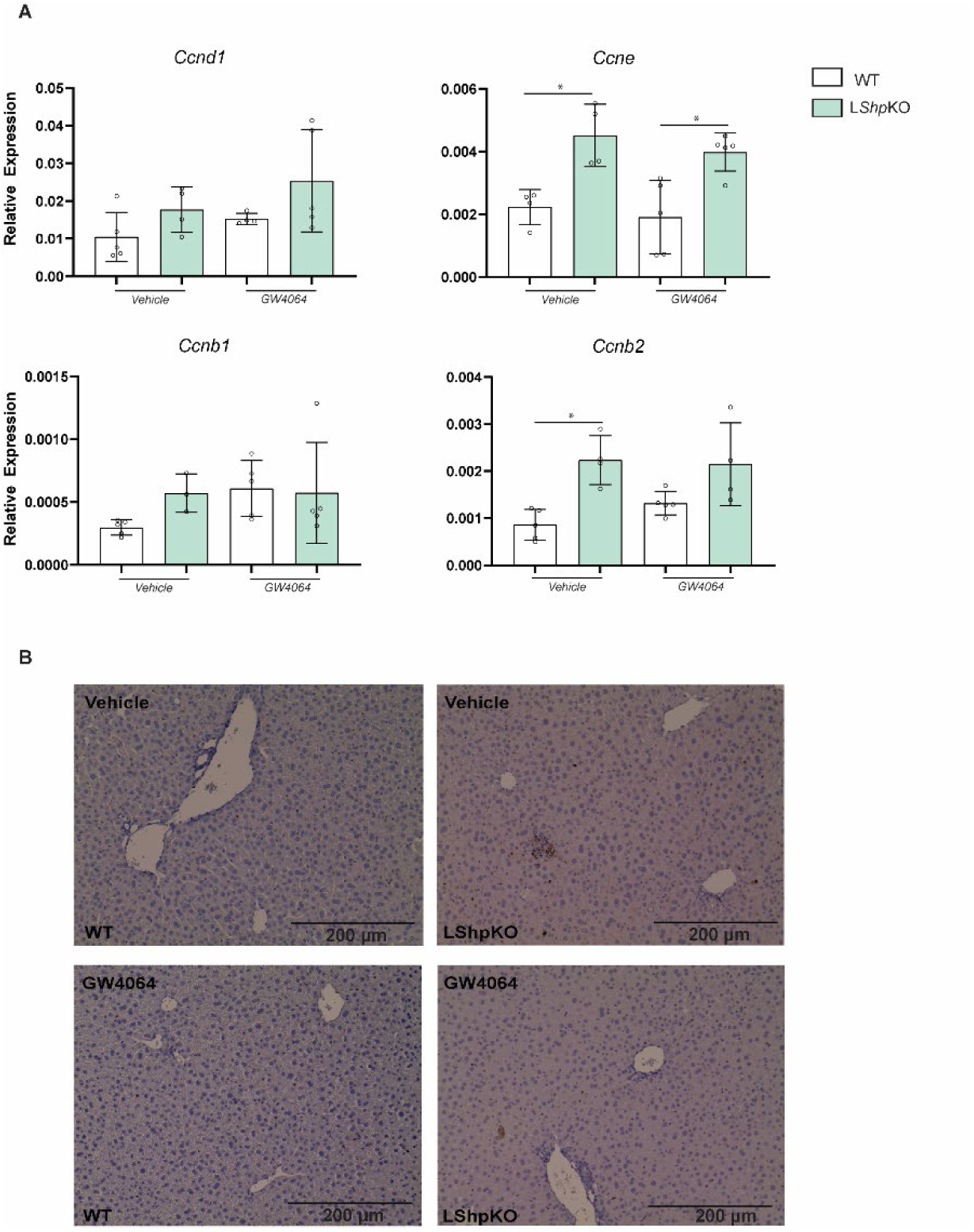
(A) Expression of cyclins expressed during the cell cycle in WT and *LShpKO* animals treated with vehicle or GW4064. (B) KI-67 immunostaining of WT and *LShpKO* mice treated with Vehicle or GW4064. One-way ANOVA analysis of variance Bonferroni test was used to determine significance between groups. *P≤.05, **P≤.01, ***P≤.001, ****P≤.0001.

### Activation of *Fxr* does not alleviate transcriptional changes caused by *Shp* deficiency

*Shp* is primarily understood as a downstream target of *Fxr* in the liver and intestine (42–44). However, *Shp* can act co-ordinately with *Fxr* to maintain bile acid homeostasis (45–48). To examine the response to *Fxr* activation in the absence of *Shp*, we treated control and *LShpKO* animals with vehicle (1% methylcellulose with 1% Triton X-100) or synthetic agonist for *Fxr*, GW4064 (50 mg/kg). The alternative BA synthesis genes *Cyp27a1* and *Cyp7b1* remained unaltered between all four groups. The canalicular export (*Mdr2*) and hepatic bile acid uptake (*Ntcp*) were increased in the absence of *Shp,* but this increase was noted irrespective of FXR activation (Figure 6 C). Intriguingly, the increase in *Cyp7a1* gene was significantly reduced and *Cyp8b1* was also lowered in *LShpKO* livers upon GW4064 treatment (Figure 6 B).

**Figure 6:**
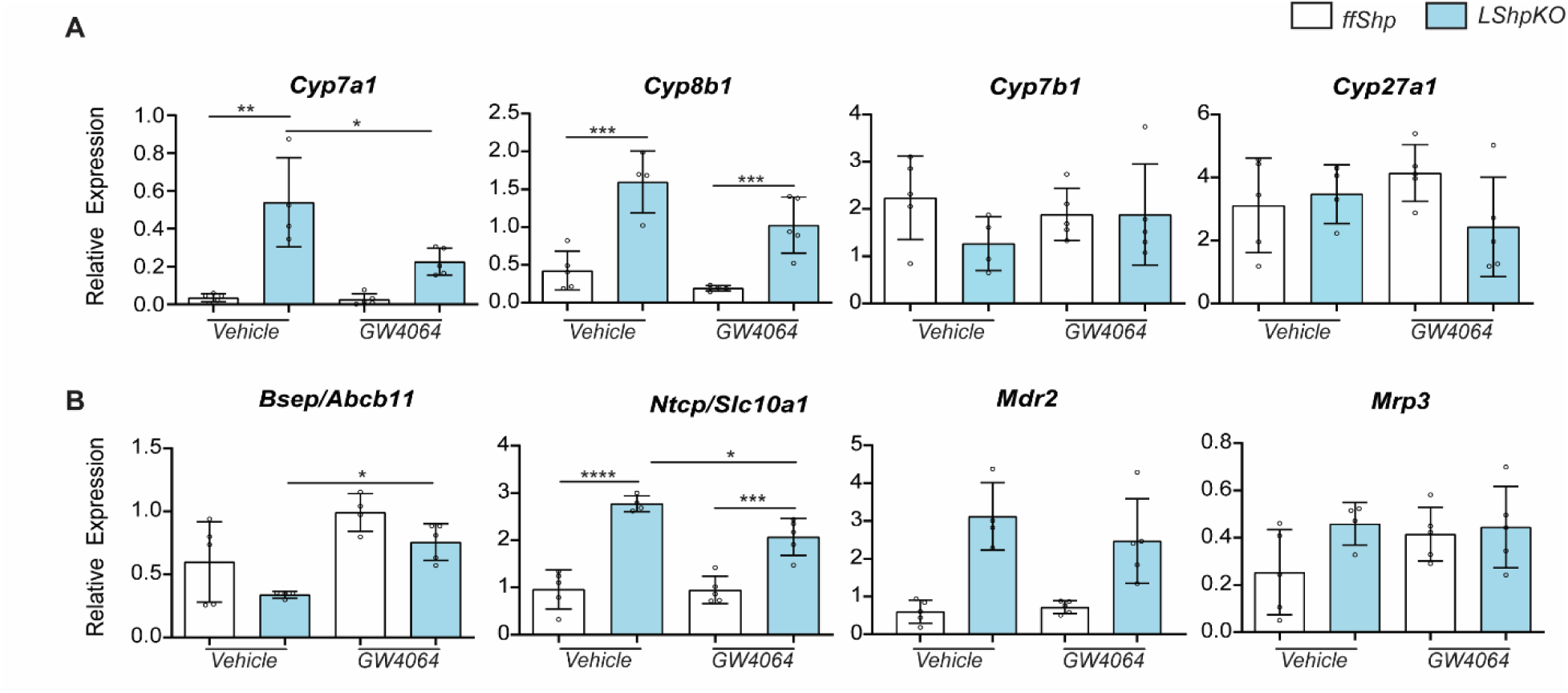
(A) Expression of key bile acid synthesis genes in WT and *LShpKO* animals treated with either vehicle or GW4064. (B) Relative expression of hepatic uptake and canalicular bile acid pumps in WT and *LShpKO* animals treated with either vehicle or GW4064. One-way ANOVA analysis of variance Bonferroni test was used to determine significance between 2 groups under 3 conditions. *P≤.05, **P≤.01, ***P≤.001, ****P≤.0001.

Next, we examined the expression profile of cyclins in *LShpKO* mice and found them unchanged post GW4064 treatment (Figure 5).

Finally, to determine genes that *Shp* uniquely regulates, we mined the previously published data (16) and identified several unique genes whose expressions are explicitly altered in the absence of *Shp* and compared them to those that are changed in the absence of *Fxr* (Supplemental Table 1 & 2) (32).

Gene ontology analysis of these unique genes revealed enrichment of circadian, amino and carboxylic acid metabolism, copper ion transport, and DNA synthesis pathways in *Shp*KO livers. In *Fxr*KO livers, we noted changes in lipid, triglyceride and nucleotide-sugar metabolism, iron ion, acetylation, chemotaxis, cholesterol, and glucose metabolism. Next, we examined if these unique genes exhibit common or overlapping motifs near their transcription start site to test whether there are new TF interacting partners of *Shp*. Using in-silico methods such as MEME-suite (49), we identified potential motifs for THAP11, ZNF143, and the motif for the Ets family of transcription factors, including ERG, ETS1, and GABPA on the promoters of genes altered in *Shp*KO livers. Because THAP11 and ZNF143 are transcription factors known to assist E2F target genes involved with cell cycle progression (50–51), we examined genes that regulate and are associated with the cell cycle. To determine if *Fxr* activation can rescue or influence the expression of these genes, we compared *LShpKO* and *ff Shp* mice treated with vehicle or GW4064 or 1%CA diet. Under basal conditions, we noted that the cell cycle genes did not change between *ff Shp* and *LShpKO* animals. Intriguingly, we found that GW-specific suppression of *E2f1*, *Acyl,* and *Cdk6* mRNA was not seen in the CA-treated samples (Figure 7 A & B). Importantly, these suppressive effects were *Shp* independent. Further, *Ezh2* transcript expression significantly decreases in the *LShpKO* mice treated with CA and shows a lower trend with GW4064 treatment.

**Figure 7:**
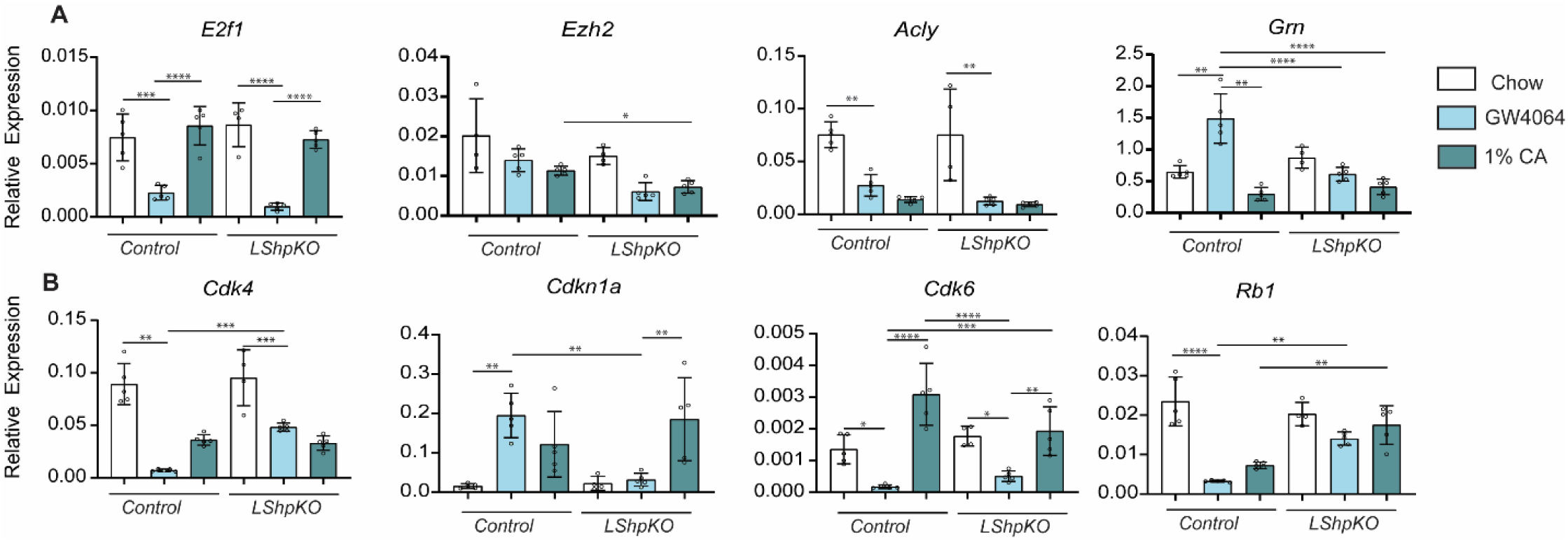
(A) Relative gene expression of several genes uniquely altered in *Shp*KO at 3 Weeks during development pertaining to liver proliferation and maturation. (B) One-way ANOVA analysis of variance Bonferroni test was used to determine significance between 2 groups under 3 conditions. *P≤.05, **P≤.01, ***P≤.001, ****P≤.0001.

Similarly, *Grn* and *Cdkn1a* gene expression is induced in a *Shp* dependent manner in WT mice treated with GW4064 but not in the CA-fed mice. This indicates a positive role for *Shp* in controlling *Ezh2 , Grn* and *Cdkn1a* transcript levels. *Rb1* transcript was strikingly decreased upon GW4064 and CA diet. But in the absence of hepatic *Shp*, CA-diet induced while GW4064 did not alter *Rb1* levels indicating a bile acid-specific response. These findings validate the role of hepatic *Shp* as a transcriptional modulator of cell cycle and proliferation both in a positive and negative direction.

## DISCUSSION

Several studies based on the global knockout of nuclear receptor *Shp* have been published over the last decades (5,6,8,14,17). Still, there is a lack of tissue-specific characterization of *Shp*. Here, we found that deletion of hepatic *Shp* leads to increased ileal bile acid concentrations and induced proliferation under chow and 1% CA conditions. We also identified several genes that are uniquely regulated by *Shp* independently of *Fxr* activation. To determine if the evolution of the SHP, FXR and BA synthesis was convergent, we conducted a phylogenetic analysis across 80 different species. Our data implicate either SHP coevolved or evolved later than FXR, as the structure of the FXR tree had two very distinct clusters of organisms formed when using the sequences with the highest percent identity. The design of the SHP tree was less distinct and more homogenous when compared to the FXR tree. Examination of the CYP7A1 and CYP8B1 trees revealed that FXR more closely resembles CYP7A1, while SHP is more similar to CYP8B1. This similarity becomes more evident as the response in the bile acid synthesis pathways is much more robust in CYP8B1 than it is in CYP7A1 (Figure 1, Figure 6).

We examined BA homeostasis in *LShpKO* mice and found that the negative feedback loop initiated by excessive amounts of bile acids is impaired. *Cyp7a1* and *Cyp8b1* genes were expressed at increased levels in the absence of *Shp* (Figure 1). In line with this data, *LShpKO* animals displayed higher bile acid content when compared to controls animals (Figure 2). To understand the increased ileal contribution to the bile acid pool in *LShpKO* mice, we analyzed ileal and hepatic transporter gene expression and did not observe significant change between control and *LShpKO* mice in these transporters (Supplemental Figure 4) (32). It is possible that their protein expression maybe altered, leading to changes in enterohepatic recirculation either due to increased hepatic secretion or decreased reabsorption, leading to higher Ileal BAs.

SHP is known to function as a transcriptional repressor and can generally be induced through various factors such as xenobiotics, genetic defects, CpG methylation, and bile acids due to their detergent chemistry (21, 52). *LShpKO* animals display increased proliferation under basal conditions (Figure 4), which was further enhanced upon CA feeding, suggesting that *Shp* could be playing a role in BA-mediated proliferation in hepatocytes and NPCs. Prolonged exposure to the CA diet in *LShpKO* animals could potentially stimulate tumorigenesis, and this data could further solidify the idea that *Shp* may be a tumor suppressor gene. We further explored the anti-proliferative role of *Shp*. Since SHP is known to repress another NR, CAR known for liver proliferation, we examined if *LShpKO* mice exhibit stark increases in cyclin genes upon CAR activation. Instead, we found a nearly complete loss of cyclin D1 upregulation in response to TC in *LShpKO* mice coupled with a compensatory increase in cyclin B2 expression. Several compensatory mechanisms are shown to exist between the cyclin genes (53) and thus may explain the lack of an exacerbated proliferative response in TC-treated *LShpKO* mice.

To identify *Shp*-specific regulation, we mined the previously published transcriptomic data from the livers of the whole body *Shp* and *Fxr* knockout mice. We identified several distinct genes represented in different signaling pathways including copper ion transport, circadian regulation and metabolism. GW4064 treatment in WT and *LShpKO* mice led us to examine the consequence of FXR activation in the presence and absence of SHP. Next, we tested and found that activation of FXR with GW4064 did not alter genes involved in BA metabolism in *LShpKO* animals. For instance, *Cdkn1a* transcript was induced upon FXR activation with GW4064 in WT mice but not in *LShpKO* animals, indicating that *Shp* expression was necessary for this induction. On the other hand, GW4064 decreased the expression of *Cdk4* in the WT mice, and this reduction was blunted in *LShpKO* mice, indicating a repressive role for SHP. Also, GW4064 did not alter proliferation as assessed using KI-67. Overall, these results show that *Shp* plays a key role in modulating tissue specific BA content, overall BA homeostasis, and proliferation in the liver.

## Notes

https://data.mendeley.com/v1/datasets/zb6mmp3zm3/draft?preview=1

### Competing Interest Statement

The authors have declared no competing interest.

