## Supplemental Figures for "Liver-specific Deletion Of Small Heterodimer Partner Alters Enterohepatic Bile Acid Levels And Promotes Bile Acid-Mediated Proliferation In Male Mice"

**Supplemental Data and Figures for:**

### Hepatic deletion of Small Heterodimer Partner alters enterohepatic bile acid content and induces a mild proliferative response in Male Mice.

**Authors:**

Ryan Philip Henry Shaw[1*], Peter Kolyvas[1*], Nathanlown Dang[1], Angela Hyon[1], and Sayeepriyadarshini Anakk[1, 2,,3#]

**Affiliations:**

[1]Department of Molecular and Integrative Physiology, [2]Beckman Institute for Advanced Science and Technology, [3]Division of Nutritional Sciences, University of Illinois at Urbana-Champaign, Urbana, IL.

* These authors contributed equally to this work.

[#] Corresponding: (SA.)

#### Supplemental Figures Legends

**Supplemental Figure 1:** Generation of *LShpKO* mice and relative mRNA expression of Major Nuclear Receptors *Fxr* and *Shp* A.) DNA gel showing PCR products of *Shp* and B) albumin cre. C) Relative mRNA expression of *Fxr-alpha* D) Relative mRNA expression of *Shp.* N=3-18 mice per group. Values shown as mean ± SD. One-way ANOVA analysis of variance Bonferroni test was used to determine significance between groups. *P≤.05, **P≤.01, ***P≤.001, ****P≤.0001.

**Supplemental Figure 2:** Relative expression of synthesis genes in chow fed mice A) *Cyp7a1* B) *Cyp8b1* C) *Cyp27a1* D) *Cyp7b1.* Relative expression of transport genes under chow diet E) *Mdr2* F) *Mrp4* G) *Mrp3* H) *Ntcp*. N=3-12 mice per group. Values shown as mean ± SD. One-way ANOVA analysis of variance Bonferroni test was used to determine significance between groups. *P≤.05, **P≤.01, ***P≤.001, ****P≤.0001.

**Supplemental Figure 3:** Relative expression of synthesis genes in 1%CA fed mice A) *Cyp7a1* B) *Cyp8b1* C) *Cyp27a1* D) *Cyp7b1.* Relative expression of transport genes under chow diet E) *Mdr2* F) *Mrp4* G) *Mrp3* H) *Ntcp*. N=4-18 mice per group. Values shown as mean ± SD. One-way ANOVA analysis of variance Bonferroni test was used to determine significance between groups. *P≤.05, **P≤.01, ***P≤.001, ****P≤.0001.

**Supplemental Figure 4:** Relative expression of transporters in the ileum of *ff Shp* and L*Shp*KO mice under chow (A-D) and 1% CA conditions (E-H). N= 5-6 mice per group. Values shown as mean ± SD. One-way ANOVA analysis of variance Bonferroni test was used to determine significance between groups. *P≤.05, **P≤.01, ***P≤.001, ****P≤.0001.

**Supplemental Figure 5:** Characterization of *Car* mediated proliferation in L*Shp*KO mice. (A) Relative expression of several cyclins involved in the progression of the cell cycle. (B) Ki-67 immunohistochemistry of liver tissue samples showing proliferation of hepatocytes and Non-Parenchymal Cells (NPCs) treated with TC. (C) Quantification of proliferating hepatocytes. Quantification of proliferating NPCs. One-way ANOVA analysis of variance Bonferroni test was used to determine significance between 2 groups under 2 conditions. Two-tailed unpaired t test was used to determine significance between 2 groups. *P≤.05, **P≤.01, ***P≤.001, ****P≤.0001.

**Supplemental Table 1:** Unique gene changes in 3 Week SHPKO mice when compared to WT mice.

**Supplemental Table 2:** Unique gene changes in 3 Week FXRKO mice when compared to WT mice.

**Supplemental Table 3:** Motifs predicted to be commonly found in genes dysregulated by the loss of *Shp* and not *Fxr.*

**Supplemental Table 4:** Motifs predicted to be commonly found in genes dysregulated by the loss of *Fxr* and not *Shp*.

**Supplemental Figure 5:** Primer sequences used throughout the present study.

#### Supplemental Figures and Tables

#### Supplemental Figure 1

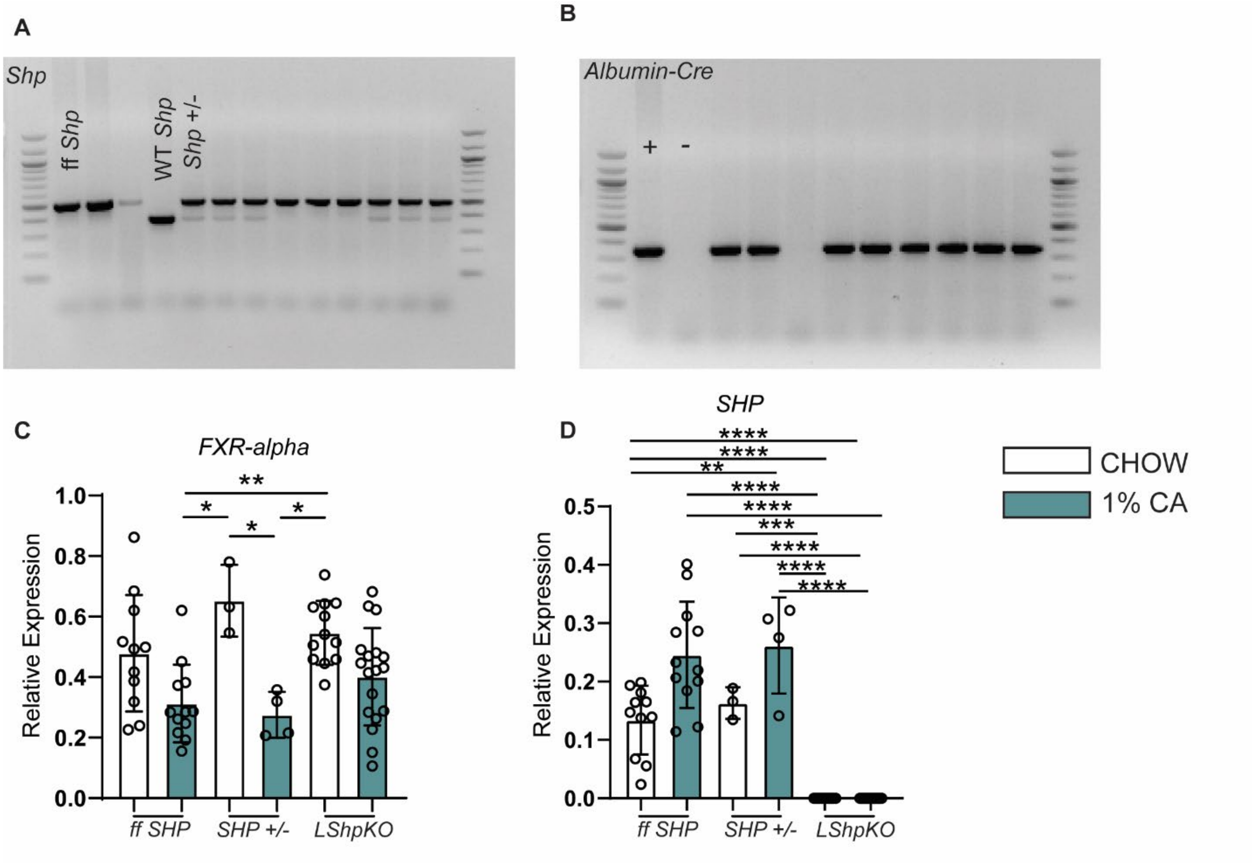

**Supplemental Figure 2**

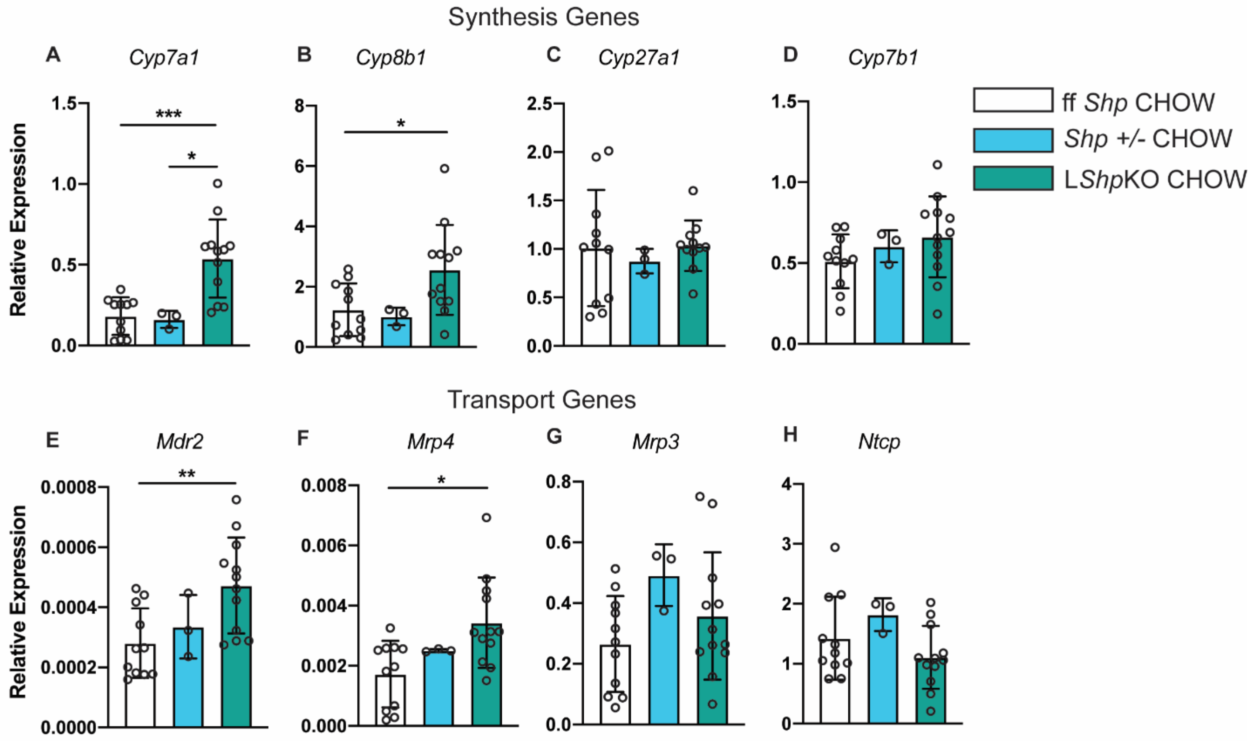

#### Supplemental Figure 3

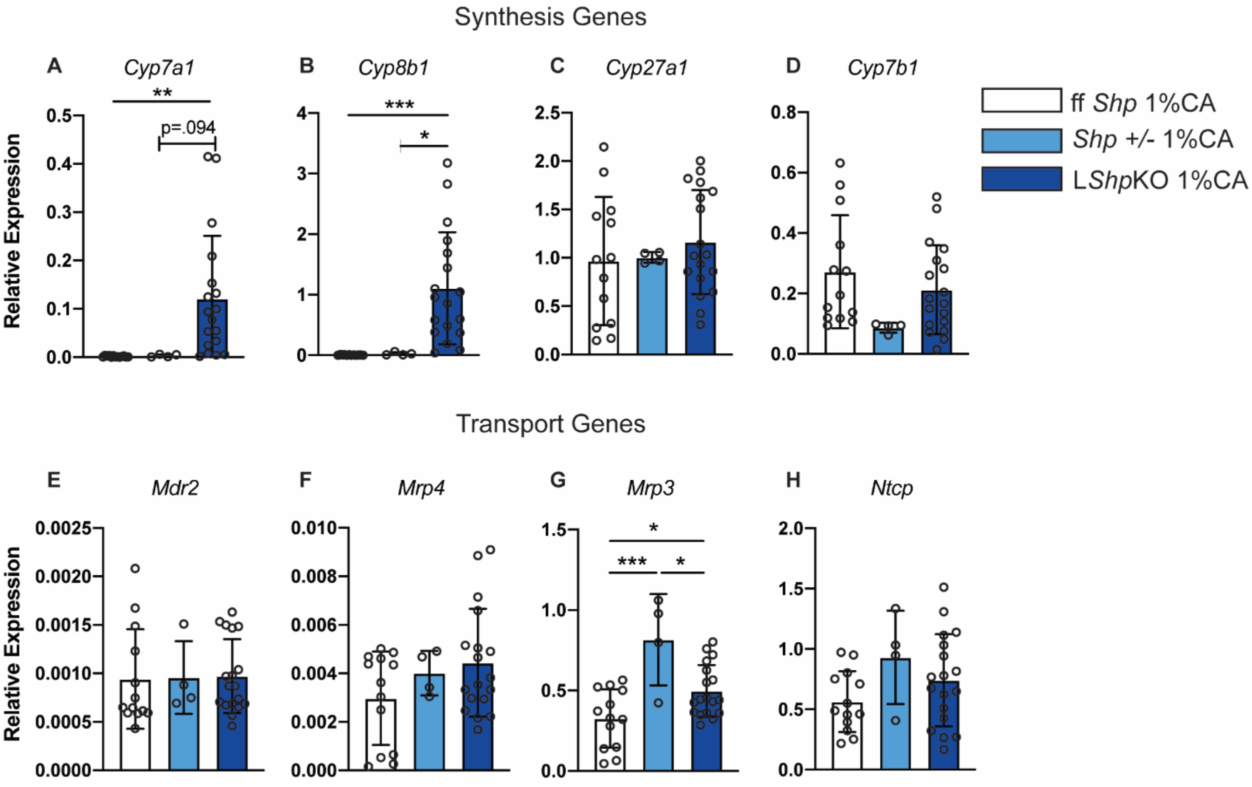

#### Supplemental Figure 4

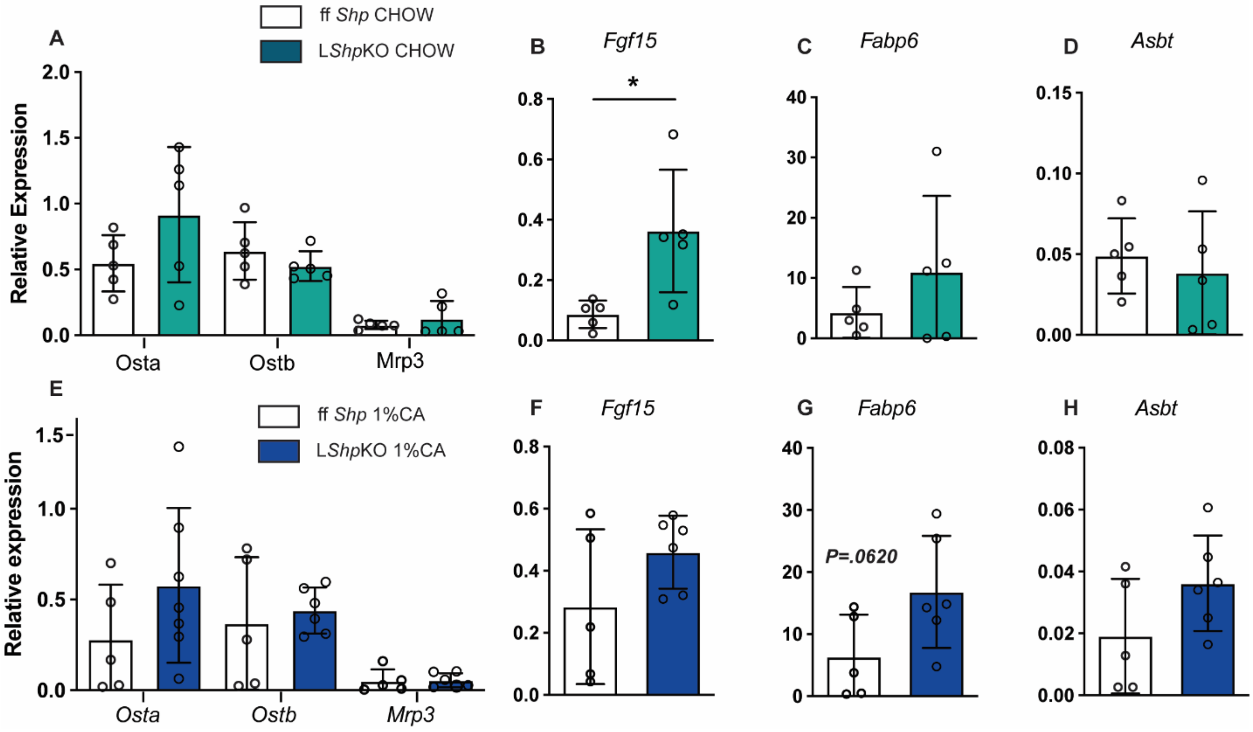

#### Supplemental Figure 5

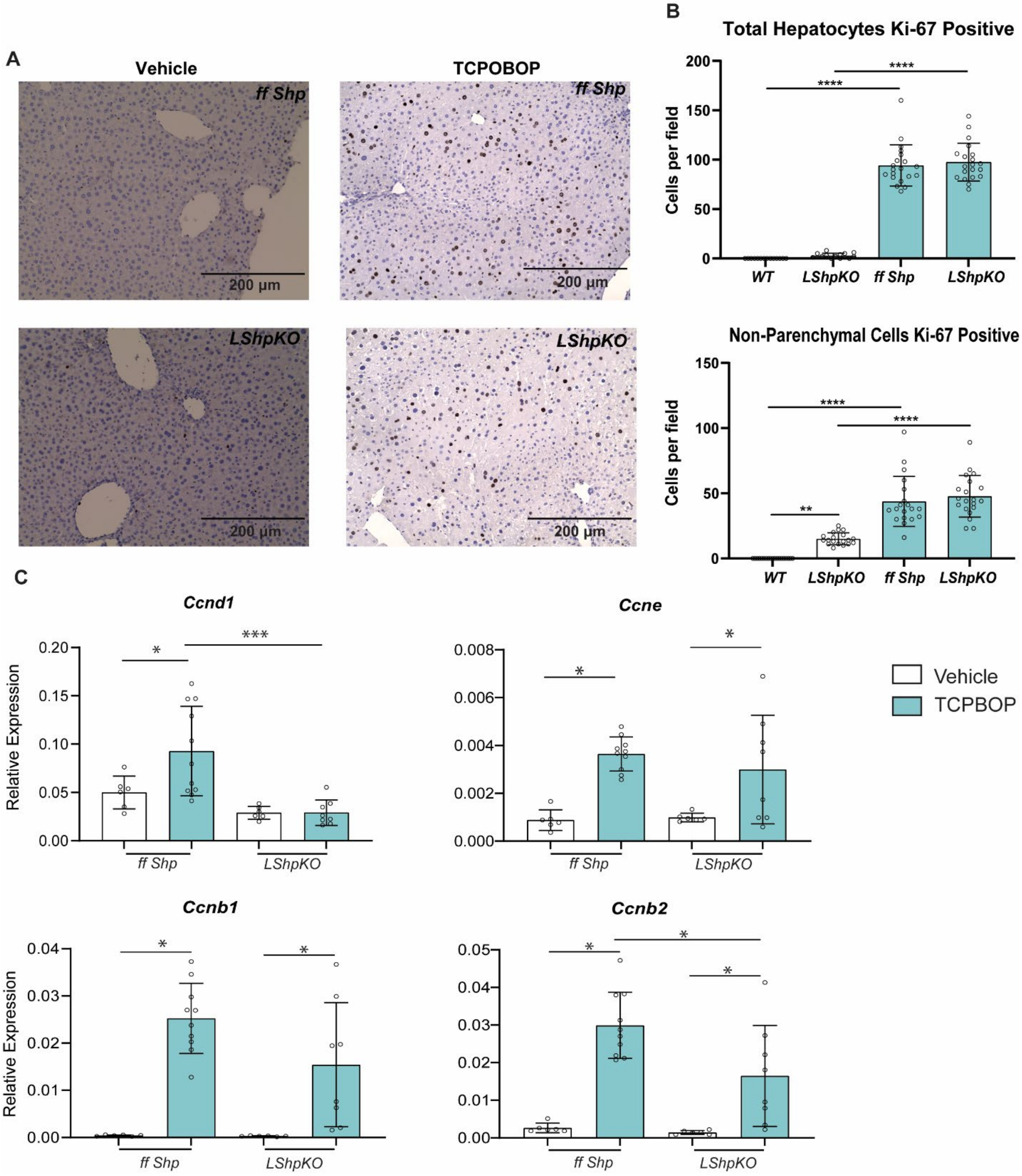

**Supplemental Table 1 (Separate Excel File)**

**Supplemental Table 2 (Separate Excel File)**

#### Supplemental Table 3

| Motif | P-value | Consensus Sequence | Sequence Logo |
|---|---|---|---|
| ZN143_MOUSE.H11MO.0.A | 6.15E-07 | KGCMTKCTGGGARWTGTAGTYY | 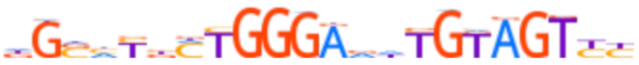 |
| THA11_MOUSE.H11MO.0.B | 2.31E-05 | KGSMTbYTGGGAvdTGTAGTYY | 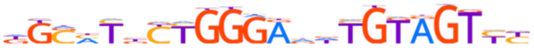 |
| GABPA_MOUSE.H11MO.0.A | 8.86E-06 | vvvvSCGGAAGYRv | 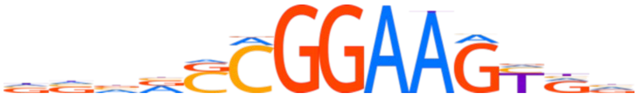 |
| ERG_MOUSE.H11MO.0.A | 5.78E-05 | vvvSMGGAARhdvv | 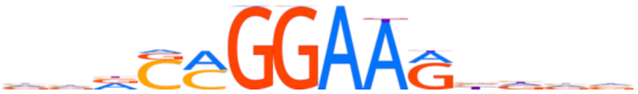 |
| ZN431_MOUSE.H11MO.0.C | 6.68E-04 | dvdbRvhRWCCTAAGACAGGvWb | 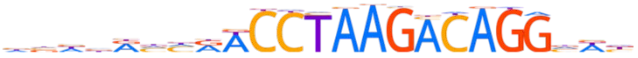 |
| ETS1_MOUSE.H11MO.0.A | 1.02E-04 | vvRSMGGAAGYRR | 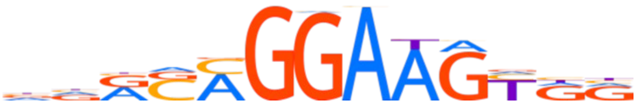 |

**Supplemental Table 4**

| Motif | P-value | Consensus Sequence | Sequence Logo |
|---|---|---|---|
| SRY_MOUSE.H11MO.0.B | 7.75E-08 | bWTTGTWWh | 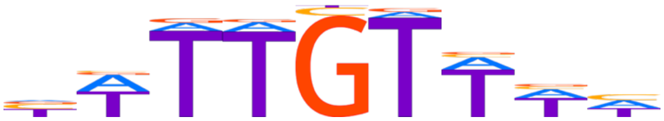 |
| OVOL1_MOUSE.H11MO.0.C | 5.88E-07 | KGTAACKGT | 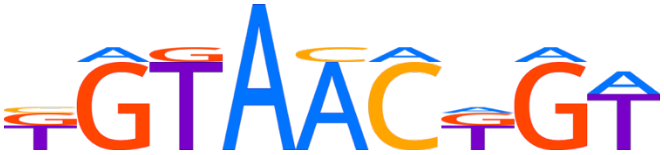 |
| PRDM1_MOUSE.H11MO.0.A | 7.52E-07 | RRRAGdGAAAGKdv | 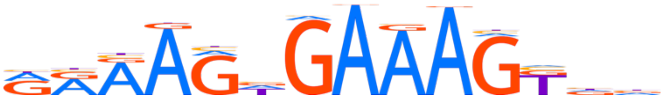 |
| CREB1_MOUSE.H11MO.0.A | 2.58E-06 | ndRTGACGYhW | 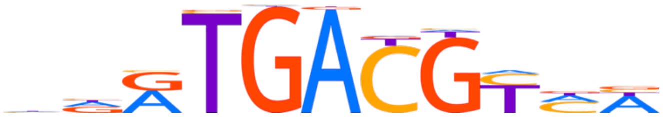 |
| LEF1_MOUSE.H11MO.0.B | 4.06E-07 | bSYTTTSWhnKbhh | 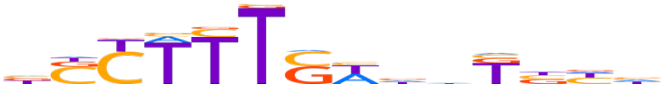 |
| FOXL2_MOUSE.H11MO.0.C | 1.32E-06 | TGTTTWYhWWh | 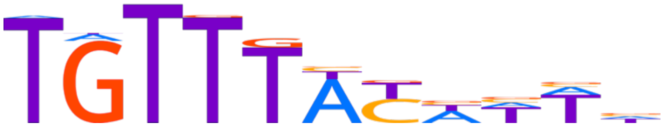 |
| ELK4_MOUSE.H11MO.0.B | 1.00E-06 | vvvSMGGAARbvv | 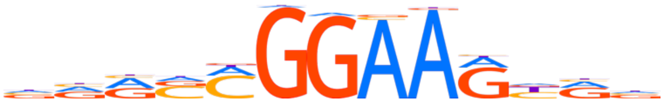 |
| NANOG_MOUSE.H11MO.0.A | 2.00E-06 | bbYWTTSWnWTKYWRAd | 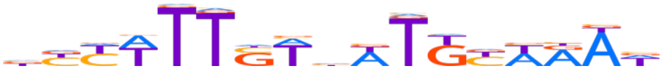 |

| | | | |
|---|---|---|---|
| PBX3_MOUSE.H11MO.0.A | 4.02E-06 | bbbYSATTGGYddv | 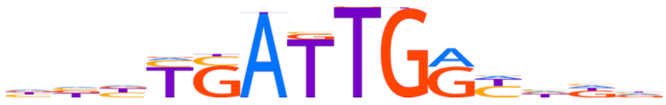 |
| FEV_MOUSE.H11MO.0.B | 1.45E-06 | vvvRGARRhS | 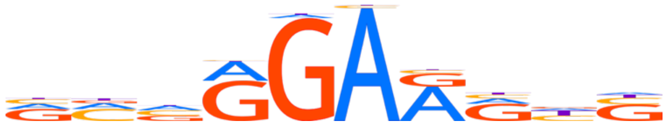 |
| FOXD3_MOUSE.H11MO.0.C | 2.32E-06 | bbTGTTTnYhYddbh | 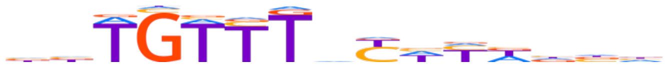 |
| NRF1_MOUSE.H11MO.0.A | 4.75E-05 | YdvYGCGCMKGCGCvvv | 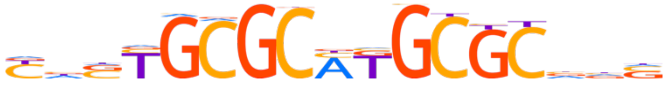 |
| CTCFL_MOUSE.H11MO.0.A | 2.80E-05 | vCCdSYAGGKGGCGChvb | 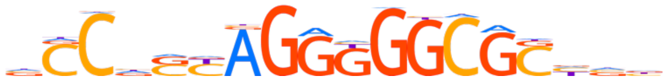 |
| TBX21_MOUSE.H11MO.0.A | 5.82E-06 | dddKGTGdSWdd | 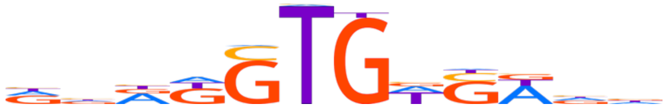 |
| IRF8_MOUSE.H11MO.0.A | 2.31E-05 | vdddvRGGAASTGAAASYdv | 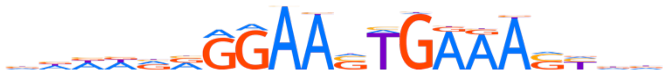 |
| CREM_MOUSE.H11MO.0.C | 4.14E-05 | SRvTGACGTSA | 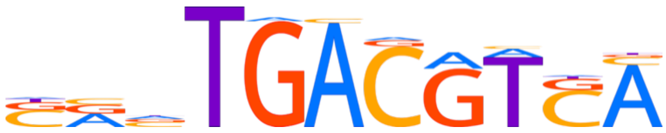 |

| | | | |
|---|---|---|---|
| TAF1_MOUSE.H11MO.0.A | 1.17E-05 | vRRRAWGGMRRhbv | 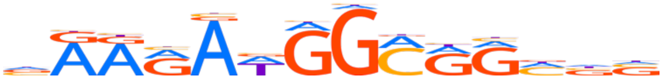 |
| ETV4_MOUSE.H11MO.0.B | 8.82E-06 | vMGGAAGY | 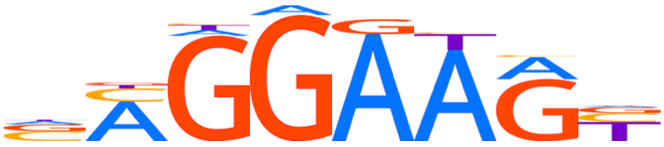 |
| MEF2A_MOUSE.H11MO.0.A | 2.14E-05 | ddYTATTTWWRKhh | 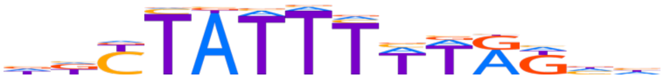 |
| MEF2D_MOUSE.H11MO.0.A | 6.35E-05 | dKYTATTTWTAKhh | 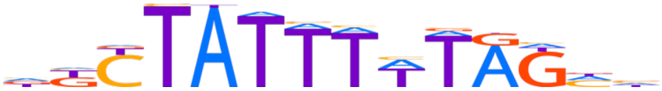 |
| FOXJ2_MOUSE.H11MO.0.C | 6.97E-05 | TRTTTATYTd | 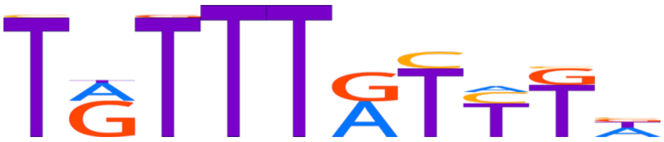 |
| ATF1_MOUSE.H11MO.0.B | 1.63E-04 | nTGACGTSMv | 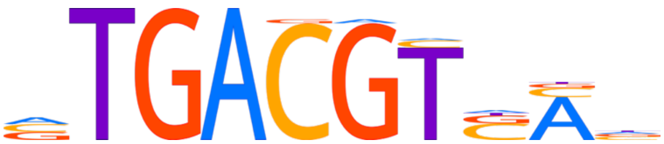 |
| FOXM1_MOUSE.H11MO.0.B | 4.75E-05 | TRTTTRYWbWbn | 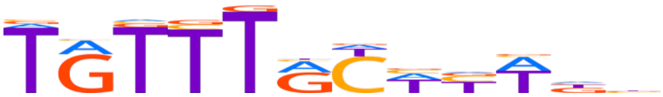 |

| | | | |
|---|---|---|---|
| ETV6_MOUSE.H11MO.0.C | 2.89E-05 | dSAGGAARb | 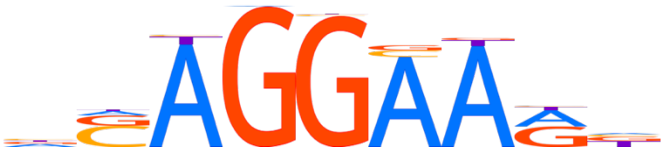 |
| E2F3_MOUSE.H11MO.0.A | 9.95E-05 | dSGCGGGARv | 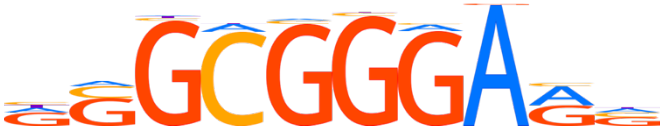 |
| SMAD3_MOUSE.H11MO.0.B | 2.99E-05 | vbYTShCWSCWb | 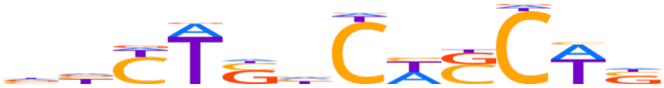 |

#### Supplemental Table 5

| Gene | NCBI Gene ID | Foward | Reverse |
|---|---|---|---|
| E2f1 | 13555 | GCCTGGAGCAAGAAGCAGT | CAGTGGTGACAGTTGGTCCT |
| Ezh2 | 14056 | GACTGCTTCCTACATCCCTTC | CTTAGCTCCCTCCAGATGCT |
| Cdkn1a | 12575 | TATCCAGACATTCAGAGCCACAG | ACTTTGCTCCTGTGCGGAAC |
| Cdk4 | 12567 | AGCCGAGCGTAAGATCCC | ACACCAATTTCAGCCACGGG |
| Cdk6 | 12571 | CCCCAGCAACCTCTCCTTC | GCCCACAATCTCTGCACTTTT |
| Rb1 | 19645 | TGCATGGCTTTCAGATTCACC | GCTGAGAGGACAAGCAGGTT |
| Acly | 104112 | GAGTCCCGAGCTGATGAAGT | GACTTGGGACTGAATCTTGGGG |
| Grn | 14824 | TGCCCGTTCTCTAAGGGTGT | ACAGCACCCAAGTTATC |
| Cyp7b1 | 13123 | TCTCTTTGCCGCCACCTTAC | TTTCAGGCCTGCCAAGAT |
| Mdr2 | 18670 | TCTTGAGGCAGCGAGAAACG | TTTCTCTGCCTTTGCTGA |
| Cyp27a1 | 104086 | CGGGGACCGGAACGCT | TTGGTCTTGTTCAGCACCTGGA |
| Cyp7a1 | 13122 | CAGGGAGATGCTCTGTGTTCA | AGGCATACATCCCTTCCGTGA |
| Cyp8b1 | 13124 | AAGGCTGGCTTCCTGAGCTT | AACAGCTCATCGGCCTCATC |
| Mrp4 | 239273 | CATACCATTGGTTCCGCTCT | TGCATCAAACAGCTCCTGAC |
| Mrp3 | 76408 | ATCATTGTGCTTGCTGGTG | CTCCTGGTCTTCATCTGGTG |
| Ntcp | 20493 | CTCAGCGTCATTCTGGTAGTT | CCAGAAGTGAGCCTTGATCTT |
| Asbt | 20494 | GGAACTGGCTCCAATATCCTG | GTTCCCGAGTCAACCCACAT |
| Fabp6 | 16204 | GGTCTTCCAGGAGACGTGAT | ACATTCTTTGCCAATGGTGA |
| Fgf15 | 14170 | CGCTACTCGGAGGAAGACTG | TTTGAGGGTTTCTGGTCCTG |
| Osta | 106407 | TTGTGATCAACCGATTTGT | CCACGCCTGTTTCATTACCT |
| Ostb | 330962 | ATCCTGGCAAACAGAAATCG | GGGTCTGGCAGAAAGACAAG |
| Ccnd1 | 12443 | TTCGTGGCCTCTAAGATGAAG | CCGGATAGAGTTGTCAGTGTAG |
| Ccne | 12447 | GATCGTTACATGGCATCACA | AAACTGGTGCAACTTTGGAG |
| Ccnb2 | 12442 | GCCAAGAGCCATGTGACTATC | CAGAGCTGGTACTTTGGTGTTC |
| Ccnb1 | 268697 | CTTGCAGTGAGTGACGTAGAC | CCAGTTGTCGGAGATAAGCATAG |
